# Genetic landscapes reveal how human genetic diversity aligns with geography

**DOI:** 10.1101/233486

**Authors:** Benjamin Marco Peter, Desislava Petkova, John Novembre

## Abstract

Geographic patterns in human genetic diversity carry footprints of population history^1,2^ and provide insights for genetic medicine and its application across human populations^3,4^. Summarizing and visually representing these patterns of diversity has been a persistent goal for human geneticists^5–10^, and has revealed that genetic differentiation is frequently correlated with geographic distance. However, most analytical methods to represent population structure^11–15^ do not incorporate geography directly, and it must be considered *post hoc* alongside a visual summary. Here, we use a recently developed spatially explicit method to estimate “effective migration” surfaces to visualize how human genetic diversity is geographically structured (the EEMS method^16^). The resulting surfaces are “rugged”, which indicates the relationship between genetic and geographic distance is heterogenous and distorted as a rule. Most prominently, topographic and marine features regularly align with increased genetic differentiation (e.g. the Sahara desert, Mediterranean Sea or Himalaya at large scales; the Adriatic, interisland straits in near Oceania at smaller scales). In other cases, the locations of historical migrations and boundaries of language families align with migration features. These results provide visualizations of human genetic diversity that reveal local patterns of differentiation in detail and emphasize that while genetic similarity generally decays with geographic distance, there have regularly been factors that subtly distort the underlying relationship across space observed today. The fine-scale population structure depicted here is relevant to understanding complex processes of human population history and may provide insights for geographic patterning in rare variants and heritable disease risk.

---

In many regions of the world, genetic diversity “mirrors” geography in the sense that genetic differentiation increases with geographic distance (“isolation by distance” ^17–19^); However, due to the complexities of geography and history, this relationship is not one of constant proportionality. The recently developed analysis method EEMS visualizes how the isolation-by-distance relationship varies across geographic space^16^ Specifically, it uses a model based on a local “effective migration” rate. For several reasons, the effective migration rates inferred by EEMS do not directly represent levels of gene flow^16^; however they are useful for conveying spatial population structure: populations in areas of high effective migration are genetically more similar than other populations at the same geographic distance, and conversely, low migration rates imply genetic differentiation increases rapidly with distance. In turn, a map of inferred patterns of effective migration can provide a compact visualization of spatial genetic structure for large, complex samples.

We apply EEMS on a combination of 27 existing single nucleotide polymorphism (SNP) datasets. In total, these comprise 6066 individuals from 419 locations across Eurasia and Africa (Extended Data Table 1), which we organize in seven analysis panels: an overview Afro-Eurasian panel (AEA), four continental-scale panels, and two panel of Southern African KhoeSan and Bantu speakers. For all analysis panels, the inferred EEMS surfaces are “rugged”, with numerous high and low effective migration features (Fig 1a, Fig 2) that are strongly statistically supported when compared to a uniform-migration model (Extended Data Table 2). The regions of depressed effective migration often align in long, connected stretches that are present in more than 95% of MCMC iterations. To facilitate discussion, we annotate these stretches with dashed lines and refer to them as “troughs” of effective migration (Figs. 1a, 2, Extended Data Figs. 2-4). Conversely, intermediate- and high-migration areas between troughs are referred to as corridors.

**Figure 1:**
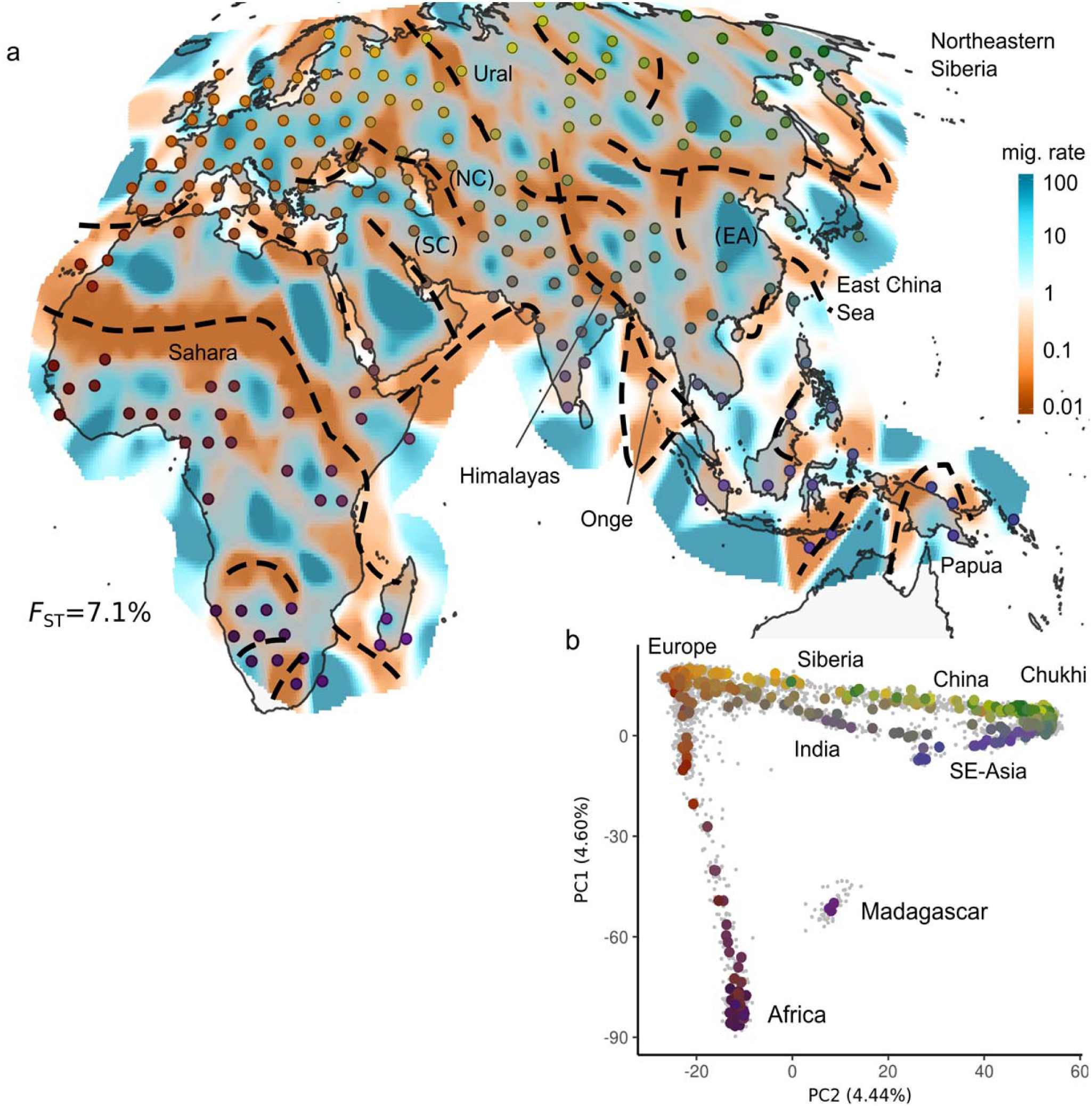
Large-scale patterns of population structure. **a:** EEMS posterior mean effective migration surface for Afro-Eurasia (AEA) panel. Regions and features discussed in the main text are labeled. Approximate location of troughs are annotated with dashed lines (see Extended Data Figure 2). **b:** PCA plot of AEA panel: Individuals are displayed as grey dots, Colored dots reflect median of sample locations; with colors reflecting geography and matching with the EEMS plot. Locations displayed in the EEMS plot reflect the position of populations after alignment to grid vertices used in the model (see methods). For exact locations, see annotated Extended Data Figure 2 and Table S1. The displayed value of *F*_*ST*_ emphasizes the low absolute level of differentiation in human SNP data.

**Figure 2:**
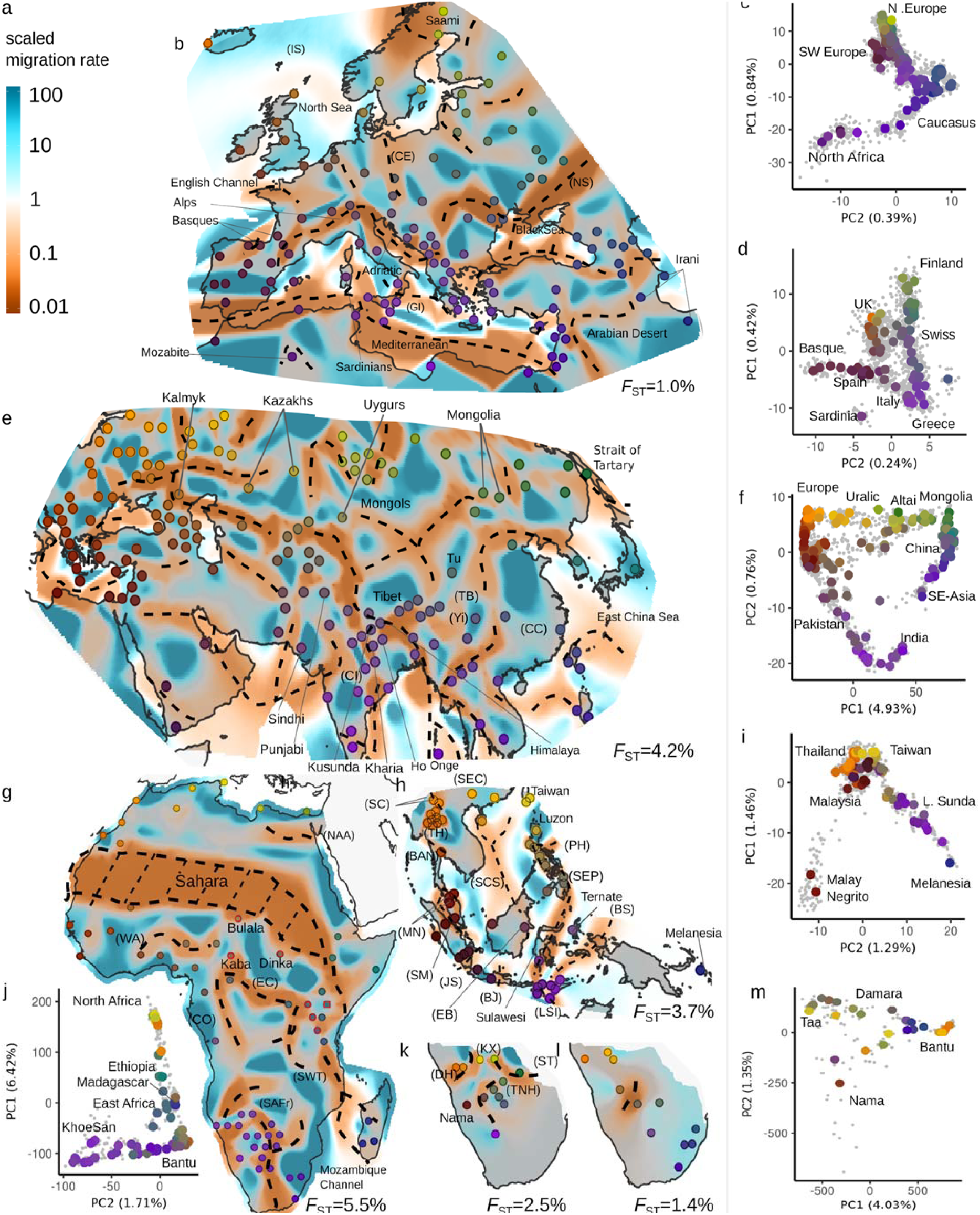
Regional patterns of genetic diversity. **a:** scale bar for relative effective migration rate. Posterior effective migration surfaces for **b**: Western Eurasia (WEA) **e**: Central/Eastern Eurasia (CEA) **g**: Africa (AFR) **h** South East Asian (SEA) **k:** Southern African KhoeSan (SAKS) **l**: Southern African Bantu (SAB) analysis panels.; In panel g, red circles indicate Nilo-Saharan speakers. Approximate location of troughs are shown with dashed lines (see Extended Data Figure 4). PCA plots: **c**: WEA d:Europeans in WEA **f**: CEA **i**: SEA **j**: AFR **m**: SAHG+SAB. Individuals are displayed as grey dots. Large dots reflect median PC position for a sample; with colors reflecting geography matched to the corresponding EEMS figure. In the EEMS plots, approximate sample locations are annotated. For exact locations, see annotated Extended Data Figure 4 and Table S1. Features discussed in the main text and supplement are labeled. *F*_*ST*_ values per panel emphasize the low absolute levels of differentiation.

In the broad overview Afro-Eurasia panel (Fig. 1; n=4,697 samples; 370 locales; *F*_*ST*_ = 0.071) we see that troughs often align with topographical obstacles to migration, such as deserts (Sahara), seas (Mediterranean, Red, Black, Caspian, East China Seas) and mountain ranges (Ural, Himalayas, Caucasus). Among the main features are several large regions that have mostly high effective migration, such as Europe, East Asia, Sub-Saharan Africa and Siberia. Several large-scale corridors are inferred that represent long-range genetic similarity, for example: India is connected by two corridors to Europe (a southern one through Anatolia and Persia ‘SC’, and a northern one through the Eurasian Steppe ‘NC’); East Asia (EA) is connected to Siberia and to southeast Asia and Oceania. The island populations of the Andaman islands (Onge) and New Guinea show troughs nearly contiguously around them – possibly reflecting a history of relative isolation ^20,21^.

Analyses on a finer geographic scale highlights subtler features (e.g. compare Europe in Fig. 1 vs Fig. 2a), and reveal that levels of differentiation differ both on both a local and continental scale (Extended Data Table 2). At these finer scales we continue to see troughs that align with landscape features, though increasingly we see troughs and corridors that coincide with contact zones of language groups and proposed areas of human migrations. For example, in Europe (Fig. 2b) we observe troughs (NS, CE) roughly between where Northern Slavic speaking peoples currently reside relative to west Germanic speakers, and relative to the linguistically complex Caucasus region. In India (Fig. 2e), troughs demarcate regions with samples of Austroasiatic and Dravidian speakers, as well as central India (CI) relative to Northwestern India (Sindhi, Punjabi) and Pakistan. In Southeast Asia (Fig. 2h), troughs align with several straits in the Malay archipelago, but we also observe a corridor from Taiwan through Luzon to the Lower Sunda Islands (LSI), and further to Melanesia, perhaps reflecting the Austronesian expansion. In Africa (Fig. 2g), a trough aligns with the Sahara desert and extends south-eastward into a geographic region that is a complex linguistic contact zone with Afro-Asiatic speakers (North), Nilo-Saharan and Niger-Congo speakers, and linguistic isolates, the Hadza and Sandawe (South). Notably, the contiguity of the South-East extension of this trough is sensitive to the inclusion of the Hadza and Sandawe (Extended Data Fig. 9). In Sub-Saharan Africa we also find corridors perhaps reflecting the Bantu expansion from West-into Southern and Eastern Africa, where contact with Nilo-Saharan speakers resulted in complex local structure. In Southern Africa, the structure seen in separate EEMS maps prepared for Bantu and Khoe-San speakers (Fig. 2k/l) appear distinct from each other, illustrating that in some cases, different language groups can maintain independent genetic structure in the same geographic region.

Different data visualization methods invariably emphasize different aspects of the data. Contrasting with EEMS, we find that the widely used, non-spatial principal component analysis (PCA) highlights large-scale geography, and the PCA-biplots typically reflect the strongest gradients of diversity in a panel. For example, PCA highlights differentiation along an Out-of-Africa differentiation axis in the AEA panel (Fig. 1b), the circum-Mediterranean and circum-Saharan distribution of diversity in Western Eurasia and Africa, respectively, and gradients from Europe into East Asia and South Asia in the Central/Eastern Eurasian panel (Fig. 2). On the other hand, EEMS emphasizes local features, in particular troughs between adjacent groups that are often imperceptible in the PCA-biplots. This is likely due to geographical information allowing EEMS to discern subtle structure from effects of uneven sampling^16^. PCA easily identifies outlier or admixed individuals (e.g. in Africa) that are not made apparent in EEMS except when exploring model fit to find populations that are fit poorly (Extended Data Fig. 8).0020Locally differentiated populations such as the Sardinians, Basques, and Finnish strongly shape the PCA results (compare Fig. 2d to e.g. ref ^17^), whereas they are typically placed in low-migration regions in EEMS. We also compare the model fit of EEMS and low-rank PCA (using the first 2, 10 and 100 components) to the observed genetic distances as a means of assessing how well each low-dimensional approach conveys structure in full genetic data. EEMS performs better for small-scale panels, but PCA provides a better fit on the larger-scale AEA and CEA panels (Extended Data Figure 5). We hypothesize EEMS tends to represent local genetic differences relatively well, and this is supported by an analysis where we stratify the residuals of genetic distances (Extended Data Fig. 6): In most panels EEMS fits best in the lowest percentiles (corresponding to local differences), and the fit quality tends to decrease for larger genetic distances.

Overall, the maps we present provide a compact summary of the complex relationship of genes and geography in human populations. In contrast to methods that identify short bursts of gene flow (“admixture”) between diverged populations^22–24^, EEMS models local migration between nearby groups to represent heterogeneous isolation-by-distance patterns. This leads to the first of a few limitations that must be considered in interpretation. In some cases, isolation-by-distance may not be the most appropriate model, and human population may overlap spatially while maintaining differentiation. This can happen either for large perioids of time (e.g. Southern Africa Bantu vs. KhoeSan speakers) or due to recent migration, displacement or admixture events (e.g. in Central Asia). In cases of population structure due to “outliers”, we found that running EEMS at the highest resolutions that are computationally feasible results in easier interpretable plots as the degrees of freedoms of the surface are high enough that these samples can be placed in regions of isolation.

Second, the maps inferred here represent a model of gene flow that predicts genetic diversity in humans sampled today – a fuller representation would represent genetic structure dynamically through time. This is especially relevant as ancient DNA data have recently suggested human population structure can be surprisingly dynamic (e.g. ref. ^25^).

Third, the effective migration rates and their scales needs be interpreted with care. In each of our maps the overall levels of differentiation are consistently low across all populations, and EEMS draws attention to where differentiation is slightly elevated or depressed relative to expectations from geographic distance. Low effective migration between a pair of populations does not imply an absence of migration nor large levels of absolute differentiation; conversely, high levels of effective migration do not imply ongoing gene flow. The emergence of migration features in the EEMS maps that align with known topography, past historical migrations, and/or linguistic/cultural distributions does not prove a causal connection and does not constitute a formal statistical test. Formally testing the influence of specific features and environmental variables on migration rates remain important future tasks that will require extending EEMS or using different frameworks^26^.

Finally, ascertainment decisions of which samples to include will affect the outcome of any analysis. When there is a feature inferred in a region with few samples, the exact positioning of the inferred change on the map will be imprecise (e.g. the trough presumably associated with the English Channel in Fig 2b). The maps of posterior variance (Extended Data Figures 2 and 4) partly convey where there is uncertainty in positioning, but caution is still warranted as the modelling assumptions will introduce further uncertainty. In other cases, the presence or absence of a particular group may open or close corridors, sometime depending on resolution. Examples of this are the Kusundas, a Nepali people with both Tibetan and Indian ancestry causing a corridor through the Himalayas, the Kalmyk, a mongolian people in Southern Russia that are linked by a corridor to Mongolia in the CEA, but not the AEA panel, and this corridor disappears if the Kalmyks are excluded from the analysis and the Hadza and Sandawe, which cause inference of a trough in Eastern Africa when included.

Nonetheless, the maps presented here provide a useful representation of human genetic diversity, that complements results from geography-agnostic methods. Our results emphasize the importance of geographical features on shaping human genetic history and help describe fine-scale patterns of human genetic diversity^27^. By using recent large-scale SNP data and a novel analysis method, our work expands beyond previous studies of gene flow in humans^28–30^. Our rugged migration landscapes suggest a synthesis of the clusters versus clines paradigms for human structure^7,8,31^: By revealing both sharp and diffuse features that structure human genetic diversity, our results suggest that more continuous definitions of ancestry in human population genetics should complement models of discrete populations with admixture. As rare disease variants are commonly geographically localized^32,33^, the maps presented here may help predict regions where clustering of alleles should be expected. The maps also annotate present-day population structure that ancient DNA and historical/archaeological studies can aim to explain.

## Material and Methods

### Merging pipeline

We obtained SNP genotype data from 27 different studies (Extended Data Table 1). Processing was done using a reproducible snakemake pipeline^34^ available under http://github.com/NovembreLab/eems-merge, heavily relying on plink 1.9^35^ for handling genotypes. The sources differ in the input format and pre-processing, however in general we performed the following steps:

1. Remove all non-autosomal, non-SNP variants
2. Map SNP to forward strand of human reference genome b37 coordinates using chip manufacturer metadata files or SNP identifiers
3. Remove strand-ambiguous A/T and G/C variants

The remaining SNPs were then merged using successive plink --bmerge commands into a single master dataset with 9,003 individuals and 1.9M SNPs but a total genotyping rate of only 20.6%. 46 SNPs were removed because different studies reported different alternative alleles. We used a relationship filter of 0.6 using the “--rel-cutoff 0.6” flag in plink to remove 667 closely related individuals or duplicates. After merging, each analysis panel had missingness rates <0.5% (AEA=0.2%, WEA=0.3%, CEA=0.2%, SEA=0.5%, AFR=0.2%, SAHG=0.1%). In all panels, all SNPs passed a one-sided HWE-test (p-value< 10^−5^), with the exception of SEA, where nine (out of 7553 SNPs) failed and were excluded.

### Data Retrieval and Filtering

#### Human Origins data set^25^

Sampling location information was obtained from table S9.4 of ref. ^25^, and the data were shared by David Reich. We used the population information in the ‘vdata’ subset of all ascertainment panels, except for the analysis where we assess ascertainment bias. The utility ‘convert’ from ‘admixtools’^22^ was used to convert the data into plink format.

#### Estonian Biocentre data

The data generated by the Estonian Biocentre^36^ were provided in plink format by Mait Metspalu on 10/30/15, along with location information where it was available. This data set contained 1,282,568 SNPs. Of those, 6770 SNPs had non-unique ids and were removed.

#### HUGO Pan-Asian SNP consortium^37^

The data were downloaded on 6/24/15 from www.biotec.or.th/PASNP. Location-metadata were obtained on the same day from the map on the same website, and individuals were matched to populations using the individual identifiers. All individuals with the same tag were assigned the median of all locations from that tag. The data were first lifted onto hg19 (with 5 out of 54794 SNPs being removed), and then re-formatted into binary plink format. Due to the small size of the chip used and the low overlap with the human origins array in particular, we only consider this data in the South-East Asian panel.

#### Uniform global sample ^38^

This data were downloaded on 6/20/15 from http://jorde-lab.genetics.utah.edu/pub/affy6_xing2010/. Sampling locations were provided by Jinchuan Xing. We used version 32 of the annotation file obtained on 6/19/15 from affymetrix.com to map SNPs onto hg19, remove strand-ambiguous SNPs and to flip SNPs that were on the minus-strand.

#### POPRES data^39^

POPRES data were obtained under dbGAP study accession phs000145 to John Novembre, and we used the data as processed in ref ^17^, and only retain individuals for which all grandparents were from the same country, and labelled the Swiss sample according to self-reported language. We used version 32 of the annotation file obtained on 6/19/15 from www.affymetrix.com (Mapping250K_sp.na32.annot.csv and Mapping250K_Sty.na32.annot.csv) to filter SNPs that did not map onto hg19 and we removed strand-ambiguous AT and GC polymorphisms.

#### African data

Data from refs ^40,41^ were obtained on 04/19/17 from David Comas’ website under http://www.biologiaevolutiva.org/dcomas/?p=607. We used version 32 of the annotation file GenomeWideSNP_6.na32.annot.csv obtained on 6/19/15 from affymetrix.com to map SNPs onto hg19, remove strand-ambiguous SNPs and to flip SNPs that were on the minus-strand.

#### South-East Asian data^42^

The data were obtained on 7/14/15 from Mark Stoneking in three different source files. After merging the three different source files, SNPs not mapping to hg19 using the annotation file GenomeWideSNP_6.na32.annot.csv were removed, as were AT and GC SNPs. Sampling locations were extracted from Figure 1 of ref ^42^

#### Mediterranean Panel^43^

Data were obtained on 8/13/15 in binary plink format from http://drineas.org/Maritime_Route/RAW_DATA/PLINK_FILES/MARITIME_ROUTE.zip. Sampling location information was obtained from Supplementary Table 3 in ref. ^43^. SNPs not mapping to hg19 using the annotation file GenomeWideSNP_6.na32.annot.csv were removed, as were AT and GC SNPs.

#### Tibetan and Himalayan data

Data from refs ^44–46^ were obtained from Choongwon Jeong and Anna Di Rienzo. We used the same filtering as in the ^44^ study, but only added the samples originating from these three studies with permission from the respective authors.

### Combining Meta-information

All sources with the exception of the Estonian Biocentre data provided (approximate) sampling coordinates. However, the level of accuracy varied between sources, with some providing specific ethnicities, some (such as POPRES) only providing country information and others just providing city- or state-level information. For POPRES-derived data, and most countries, we assigned individuals to the country’s centerpoint, with the exception of Sweden and Finland, which were assigned their capital. For the Estonian Biocentre data, sampling location data were highly heterogeneous. Samples that could not be confidently assigned to a region with an accuracy of 100km were excluded. For populations with samples from multiple studies, the most accurate source location was used. For locations covered with different accuracy, only the most accurate samples were retained. For example, we dropped all Spanish individuals from POPRES (only country level data), as the Human Origins data provided higher resolution, with samples from eleven different regions in Spain. The resulting table is given as Table S1.

#### Language data

To validate troughs correlating with presumed language barriers, we cross-referenced the genetic data with linguistic data from the Glottolog 3.2 database ^47^. To do so, we compared the correlation of pairwise genetic distance and geographic distances within and between pairs of language groups. As there was frequently no primary data recording the language of speakers, we proceeded as follows: For population identifiers that correspond to languages / or ethnic groups with a clear majority language, we used that language. For samples with country-level information where the country has a clear majority language (e.g. Germany, Slovenia), that language was assigned (Table S1). Otherwise, if a sample was from a region with a clear majority language that is not obviously due to recent colonization, that language was assigned. All other samples were not assigned a language. For simplicity, we group Nilotic, Central Sudanic and Mande languages into “Nilo-Saharan”, Khoe, Kxa and Tuu speakers into “KhoeSan” and Armenic, Circassian, Kartvelian and Nakh-Daghesanian into “Caucasus”. For all troughs we hypothesize that they align with boundaries between linguistic groups, we now perform a partial mantel test comparing genetic distances and language groups as a categorical variable using the implementation in the R-package “vegan”^48^. We note that results need to be interpreted cautiously, as the mantel test is generally poorly calibrated for spatially autocorrelated data^49^.

#### Samples omitted from model fitting

Besides samples whose geographic origin we could not unambiguously assign (n=74), we removed a small number of samples that would violate some assumptions of the EEMS model. In particular, we excluded all Jewish samples (n=379), due to complexity of the diaspora and subsequent local admixture^50^) and Han-Chinese in Taiwan and Singapore(n=170), who both are recent migrant population to those locales. To avoid any possible distortion due to uneven sampling, we downsampled all single locales to at most 50 individuals, drawn independently for different panels. This resulted in a total of 6066 individuals used in at least one panel (Table S1).

### Visualization pipeline

We developed a second pipeline using snakemake^34^ to perform all subsetting and demographic analyses, available under github.com/NovembreLab/eems-around-the-world. The pipeline allows for defining panels using a flexible set of features, latitudinal and longitudinal boundaries, continent or country of samples, source study, as well as the addition and exclusion of particular samples or populations. Based on these subsets, different modules allow performing EEMS and PCA analyses, as well as generating all the figures, that were then annotated using inkscape. All configuration variables are stored in json and yaml config files. We perform EEMS and PCA for each panel independently. Structural variants are a potential confounding factor for genome-wide SNP based analysis. In PCA, these variants may result in a number of neighboring SNP in high LD to have very high loadings, thus overemphasizing the effect of these variants. For this reason, it is advisable to remove regions containing SNP that have extremely high loadings on some Principal component. Thus, for each panel, we perform a preliminary PCA analysis using flashpca^51^. The loading-scores for each PC were normalized by dividing them by the standard deviations on each PC [outlier_score = L[i]/sd(L[i])], and then we removed a 200kb window around any SNP for which |outlier_score| > 5. We also dropped individuals with more than 5% missingness, and SNPs with more than 1% missing data from each panel.

#### EEMS

To generate the map surfaces, we must choose a grid size and boundaries. Choosing a coarse grid results in faster computation, but only produces a map with broad-scale patterns. A finer grid, on the other hand, is able to reveal more details, but at a steep increase in computational cost and with an increased danger of introducing patterns that are harder to interpret. Grid density and sizes are given in Extended Data Table 1, along with population level *F*_*ST*_ calculated using plink, and *F*_*ST*_ based on the mean migration rate inferred by eems and equilibrium stepping stone model theory^52^.

We evaluated the impact of SNP ascertainment bias by running EEMS on the multiple, documented SNP ascertainment panels of the Human Origins data^25^. We found that while ascertainment bias has an effect on the heterozygosity surfaces that EEMS estimates, the migration surfaces remain relatively unaffected (Extended Data Fig. 1). Therefore, we restrict our presentation to the migration surfaces.

For each panel, we performed four pilot runs of 2-8 million iterations each. The run with the highest likelihood was then used for a second set of four runs of 4-10 million iteration each, with the first 500,000 million discarded as burn-in. Number of iteration were chosen such that total computation time was around 10 days. Every 20,000th iteration was sampled. EEMS approximates a continuous region with a triangular grid, which has to be specified. We generated global geodesic graphs at three resolutions (approximate distance between demes of 120, 240 and 500km, respectively) using dggrid v6.1^53^ and intersected these graphs with the area representing each panel (Extended Figures 2,3). All other (hyper-)parameters were kept at their default values^16^. We compared EEMS to an isolation-by-distance model with a constant migration rate by re-fitting EEMS allowing only a single migration rate tile, but arbitrary diversity rate tiles using the otherwise same settings. The resulting log Bayes Factors are given in Extended Data Table 2.

#### Evaluating fit of EEMS and PCA to genetic distances

For EEMS, the posterior samples imply an expected distance matrix between populations. For PCA, the components and their loadings provide an approximation to the genetic distance matrix between individuals. We use the median PCA values of individuals across ten PC components to produce an expected genetic distance matrix between populations. We use ten PC components as most investigators evaluate population structure based on only the first several PCs. For each method the expected genetic distance matrices are compared to the observed matrices using a simple linear correlation computed between all pairwise distances.

## Acknowledgements

We are grateful for helpful comments from Choongwon Jeong, Matthew Stephens, Anna Di Rienzo, Melinda A. Yang, Joshua G. Schraiber and members of the Novembre lab. This research was supported by research grants NIH/NCI U01 CA198933 and NIH/NIGMS R01 GM108805 (J. N. and B. M. P) and by a Swiss National Science Foundation early postdoc mobility fellowship (B. M. P.). We acknowledge the University of Chicago Research Computing Center for support of this work. We dedicate the paper in memoriam of Brad McRae (1966-2017) whose work on resistance distances underlies the EEMS methodology.

## Author contributions

B.M.P. analyzed data. B.M.P., D.P., and J.N. interpreted results. B.M.P and J.N conceived the study and wrote the manuscript.

## Supplementary Text on Regional Scale Analyses

Here we provide a more expanded discussion of the regional-scale results. To help identify features that we discuss, we have added labels to discussed features in the figures, and refer to them in the text here in parentheses. The labels are typically capitalized abbreviations and in some cases are full words.

### Western Eurasia

Europe appears largely homogeneous in the Afro-Eurasia panel, but a finer-scale analysis (Western Eurasia panel, Fig. 2a; n=2,049; 122 locales, *F*_*ST*_=0.010) reveals abundant fine-scale structure: bodies of waters are consistently covered by lower effective migration regions, with migration being lower in southern seas (Mediterranean, Adriatic, Black Sea) relative to those in northern Europe (North Sea, Irish Sea, English Channel). Terrestrial barriers are observed in: the Alps (and an adjacent region extending into Southern France), surrounding the Mozabites in Tunisia and the Saami in Scandinavia, the western and northern edges of the Arabian Desert (though we note the region has few samples). Troughs reflecting historical domains are observed: between Germanic and Northern Slavic-speakers (CE, Extended Data Figure 7) and between domains of Slavic-speakers and the Caucasus (NS).

Remaining regions are generally inferred to have average or above average migration, with one obvious corridor being that between Iceland and Scandinavia (IS), presumably due to the recent colonization of Iceland. One interesting feature is an area of East-West low migration between the Italian peninsula and Greece (GI). A corridor between Crete and Sicily is inferred south of it, and between mainland Greece and southern Italy north of it. This likely reflects a pattern of close genetic similarity among coastal Mediterranean populations observed previously^43^ but suggests it may have north-south structure. Ancient DNA results suggest that the patterns we observe are recent^54,55^ and have been shaped in the last 3,000-5,000 years with contributions from multiple sources. Strikingly, proposed expansion routes through the Eurasian Steppe and Levant into Europe partially align with corridors of high effective migration.

### Central/Eastern Eurasia

The Central/Eastern Eurasia surface (Fig. 2e; n=2,578; 181 locales, *F*_*ST*_=0.042) is overall similar to the patterns seen in the AEA panel, with a trough through the Himalayas/Tien-Shan but we observe a corridor from Mongolia to the Caspian Sea, possibly reflecting the Mongol empire, as the Kalmyk, Kazhaks, and Uygurs all have well documented Mongolian-like genetic ancestry. The presence of this corridor depends on a small number of samples; if Uygurs and Kalmyks are removed, we find a pattern similar to that in the AEA panel in that region.

Where the global analysis did not reveal any strong patterns in South Asia, at the higher resolution we observe troughs in the Indian subcontinent between a central Indian region (CI) of mainly Indo-Aryan languages and an Eastern and Southern region with two Austroasiatic speaking (Kharia, Ho), and Dravidian speaking populations, but this trough is not significantly correlated with linguistic group (Extended Data Figure 7d) We also find that the Himalayas generate a trough between India and Tibet, but the Kusunda population adds a corridor there, which is explained by the fact that they have both Tibetan and Indian ancestry ^56^.

In East Asia, we observe marine troughs in the East China Sea, strait of Tartary and the Andaman Sea (Onge). Terrestrially, we observe troughs between coastal China (CC), a central region with several Tibeto-Burman samples (TB, along with the Tu who speak a Mongolic language, and have been suggested to have received European admixture 1,200y ago^24^), and a western region anchored by Tibetan samples. The coastal Chinese region contains a high-effective-migration area that extends into Korea and Japan.

Overall, the Central/East Asia panel is particularly complex with one of the lowest levels of r^2^ between EEMS expected genetic distances and the observed distances (r^2^ = 0.66, Extended Data Fig. 5), and the residuals are very high (Extended Data Fig. 8). This is expected as the relatively open steppe has been the site of repeated long-range population movements and invasions, by e.g. Bronze Age Steppe populations, Mongols and Turkic speakers, that we expect are difficult to depict using the model of steady-state gene flow model fit by EEMS.

### South-East Asia

In the South-East Asian panel (n=1,054, 58 locales; *F*_*ST*_=0.037; Fig. 2k) troughs align with the many seas and channels in this region: the South-Chinese Sea (SCS), the waterway running east of the Philippines (PH) and Sulawesi south to the Flores Sea (SEP), the waterway between western New Guinea into the Banda Sea (BS), the Malacca strait between Sumatra and Malaysia (SM), the Sunda Strait between Java and Sumatra (JS), the Java Sea between Bali and Java (BJ), as well as the Makassar strait and Celebes Sea between Borneo and Sulawesi (EB). Two corridors, one from Taiwan/Luzon through Western Mindanao to Sulawesi, and one from Ternate through the Lower Sunda Islands (LSI) into Melanesia possibly reflect the Austronesian expansion that started roughly 3,000 years ago^57^. On the mainland, we find low effective migration north of Bangkok (BAN) and near samples from Northern Thailand (TH) (including the Southern Chinese Wa and Jinuo samples (SC)). These two samples have low inferred effective migration with South-Eastern Chinese samples (SEC). Two Malay Negrito samples in Northern Malaysia (MN) are placed in a trough, revealing their genetic distance to other South-East Asian populations also apparent on PC2 (Figure 2l).

### Africa

In Africa (AFR, n=749, 71 locales, *F*_*ST*_=0.055; Fig. 2g) two troughs corresponding to the Sahara desert and Mozambique Channel are observed. In Northern Africa (NAA), we see a small trough of low effective migration separating two latitudinal corridors; one following the Mediterranean coast and one inland (Fig 2g). The inland corridor disappears in our lower-resolution Afro-Eurasia panel (Figure 1a) and presumably reflects Sub-Saharan ancestry in some Moroccans, perhaps through trans-Saharan trade.

In the AFR-panel, we observe a trough reflecting the language group boundaries between Niger-Congo and Afro-Asiatic language speakers^58^ (Extended Data Figure 7), with the West-African Afro-Asiatic speaking Hausa and Mada being placed in a barrier together with the admixed Fulani ^59^. West Africa appears as a high-gene-flow region (WA), and two corridors pass from Nigeria - one along the coast of Congo (CO) southwards and another further east (EC) connecting to Kenya and Tanzania. In both Central and Eastern Africa Nilo-Saharan and Niger-Congolese speakers overlap, resulting in low effective migration uncorrelated with langu: the Nilo-Saharan Kaba, Dinka and Bulala are in a region of high gene flow, separated by a trough from the Biaka and Mbuti Pygmies. Southern and Eastern Africans are separated by low effective migration through Mozambique and South-Western Tanzania (SWT).

### Southern Africa

Patterns in southern Africa are complex, with a troughs separating out Western Bantu speaking populations, and the KhoeSan Khwe and Xuun. Stratifying Southern Africans into a KhoeSan (SAKS, n=109, 16 locales, FST=0.025, Fig. 2k) and Bantu speakers (SAB, n=30, 11 locales, FST=0.014; Fig. 2l) reveals very different spatial structure. The Bantu speakers are separated by a single barrier into an eastern and western location. For the South African Hunter-Gatherers most samples fall into a central region with high effective migration, including the Taa, Naro and Hoan (TNH). Troughs in the North separate this region from the Sua and Tswa (ST) and in the south-west from the Khomani and Nama (Nama), respectively. The remaining samples fall either into a Northern high migration area (Khwe and Xuun, KX) or a North-Western low migration area (Damara and Haiom, DH). These results are broadly consistent with existing work on African population structure^59–62^, and emphasize African population structure appears largely determined by the Sahara desert, the Bantu and Arabic expansions, and the complex structure of hunter-gatherer groups specifically in South Africa.

## Extended Data

**Extended Data Table 1:** Data Sources. Abbrev: Abbreviation; Ind: total number of individuals; Loc. Number of unique sample locations

| Study | Abbrev. | Samples | Locations | Source |
| --- | --- | --- | --- | --- |
| Bryc et al. 2009 | B09 | 121 | 11 | Ref <sup>40</sup> |
| Behar et al. 2010 | Be10 | 295 | 22 | Ref <sup>50</sup> |
| Behar et al. 2013 | B13 | 131 | 20 | Ref <sup>63</sup> |
| Bigham et al. 2010 | Bi10 | 45 | 3 | Ref <sup>45</sup> |
| Chaubey et al. 2011 | C11 | 37 | 5 | Ref <sup>64</sup> |
| Cardona et al. 2014 | C14 | 192 | 37 | Ref <sup>65</sup> |
| Di Cristofaro et al. 2013 | D13 | 14 | 3 | Ref <sup>66</sup> |
| Fedorova et al. 2013 | F13 | 30 | 6 | Ref <sup>67</sup> |
| HUGO Consortium 2009 | H09 | 975 | 47 | Ref <sup>37</sup> |
| Hunter-Zinck et al. 2010 | H10 | 85 | 1 | Ref <sup>41</sup> |
| Jeong et al. 2017 | J17 | 53 | 2 | Ref <sup>44</sup> |
| Kovacevic et al. 2014 | K14 | 70 | 6 | Ref <sup>68</sup> |
| Lazaridis et al. 2014 | L14 | 1590 | 159 | Ref <sup>69</sup> |
| Metspalu et al. 2011 | M11 | 127 | 11 | Ref <sup>70</sup> |
| Migliano et al. 2013 | M13 | 68 | 6 | Ref <sup>71</sup> |
| Nelson et al. 2008 | N08 | 531 | 29 | Ref <sup>39</sup> |
| Paschou et al. 2014 | Pa14 | 626 | 29 | Ref <sup>43</sup> |
| Pierron et al. 2014 | Pi14 | 114 | 5 | Ref <sup>72</sup> |
| Raghavan et al. 2014 | R14 | 83 | 9 | Ref <sup>73</sup> |
| Rasmussen et al. 2010 | R10 | 101 | 9 | Ref <sup>74</sup> |
| Rasmussen et al. 2011 | Ra11 | 19 | 3 | Ref <sup>56</sup> |
| Reich et al. 2011 | Re11 | 106 | 16 | Ref <sup>75</sup> |
| Skoglund et al. 2014 | S14 | 15 | 1 | Ref <sup>76</sup> |
| Xing et al. 2010 | X10 | 92 | 4 | Ref <sup>38</sup> |
| Xu et al. 2011 | X11 | 28 | 3 | Ref <sup>46</sup> |
| Yunusbayev et al. 2012 | Y12 | 183 | 14 | Ref <sup>77</sup> |
| Yunusbayev et al. 2015 | Y15 | 299 | 42 | Ref <sup>36</sup> |

**Extended Data Table 2:** Analysis Panels. Abb. Panel Abbreviation. Res. Avg. distance between grid points (in km); Support: log Bayes factor in favor of complex vs constant migration model. Implied F_ST_ between adjacent demes based on posterior mean migration rates. Equation 19a from ^52^ is used to calculate implied F_ST_ using a torus approximation: For F_ST_ (adjacent demes): *F*_*ST*_*=(1+32m/S(d))*^*-*1^ where S(d) is a function of the distance between demes and given by equation A12 in ^52^. In the first column, we use S(1), in the second S(4) for highest and S(2) for medium resolution panels to get FST for demes at the lowest resolution (∼500km).

| Panel | Abb. | Ind. | Locations | SNPs | Grid Size<br>(# of demes) | Resolution<br>(km) | $F_{ST}$ | Model-fit $F_{ST}$<br>(adjacent<br>demes) | Model-fit $F_{ST}$<br>(500km) | Support<br>(log-BF) |
| --- | --- | --- | --- | --- | --- | --- | --- | --- | --- | --- |
| Afro-Eurasia | AEA | 4697 | 370 | 19972 | 686 | 500 | 0.071 | 0.99% | 0.99% | 254,472 |
| Central/Eastern<br>Eurasia | CEA | 2578 | 181 | 21045 | 1147 | 240 | 0.042 | 0.22% | 0.42% | 129,035 |
| Western Eurasia | WEA | 2049 | 122 | 26438 | 1437 | 120 | 0.010 | 0.75% | 1.08% | 46,210 |
| South-East Asia | SEA | 1054 | 58 | 7553 | 1388 | 120 | 0.037 | 0.29% | 0.56% | 13,654 |
| Africa | AFR | 749 | 71 | 20984 | 694 | 240 | 0.055 | 0.81% | 1.18% | 51,771 |
| Southern Africa<br>KhoeSan | SAKS | 109 | 16 | 532343 | 227 | 120 | 0.025 | 0.32% | 0.62% | 2298 |
| Southern Africa<br>Bantu | SAB | 30 | 11 | 65095 | 227 | 120 | 0.014 | 0.26% | 0.56% | 126 |

**Extended Data Figure 1:**
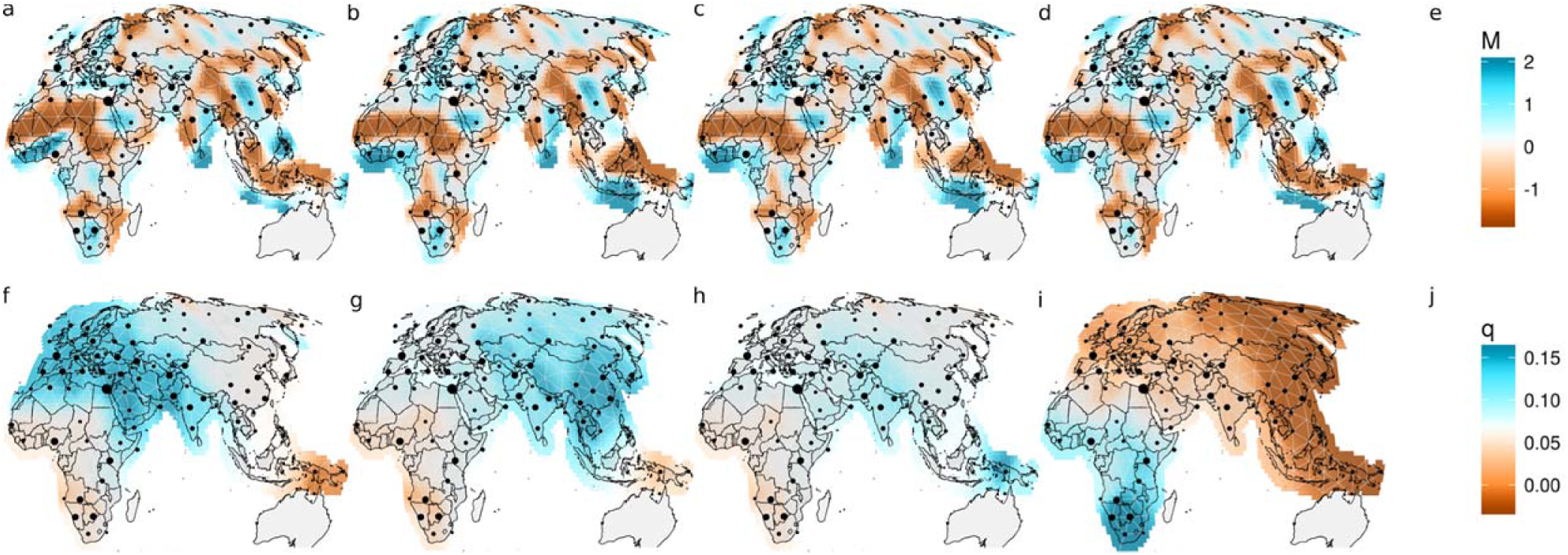
Ascertainment bias. We run EEMS only using the Human Origin data ^25^, using SNPs ascertained in a French (a/f), Chinese (b/g), Papuan (c/h) and San(d/i) individual. Migration rate surfaces (a-d) remain robust, whereas the within-deme diversity surfaces (f-i) show highests diversity at the respective ascertainment location. e/j: scale bars for migration rates and within-deme diversity rate parameters, respectively.

**Extended Data Figure 2:**
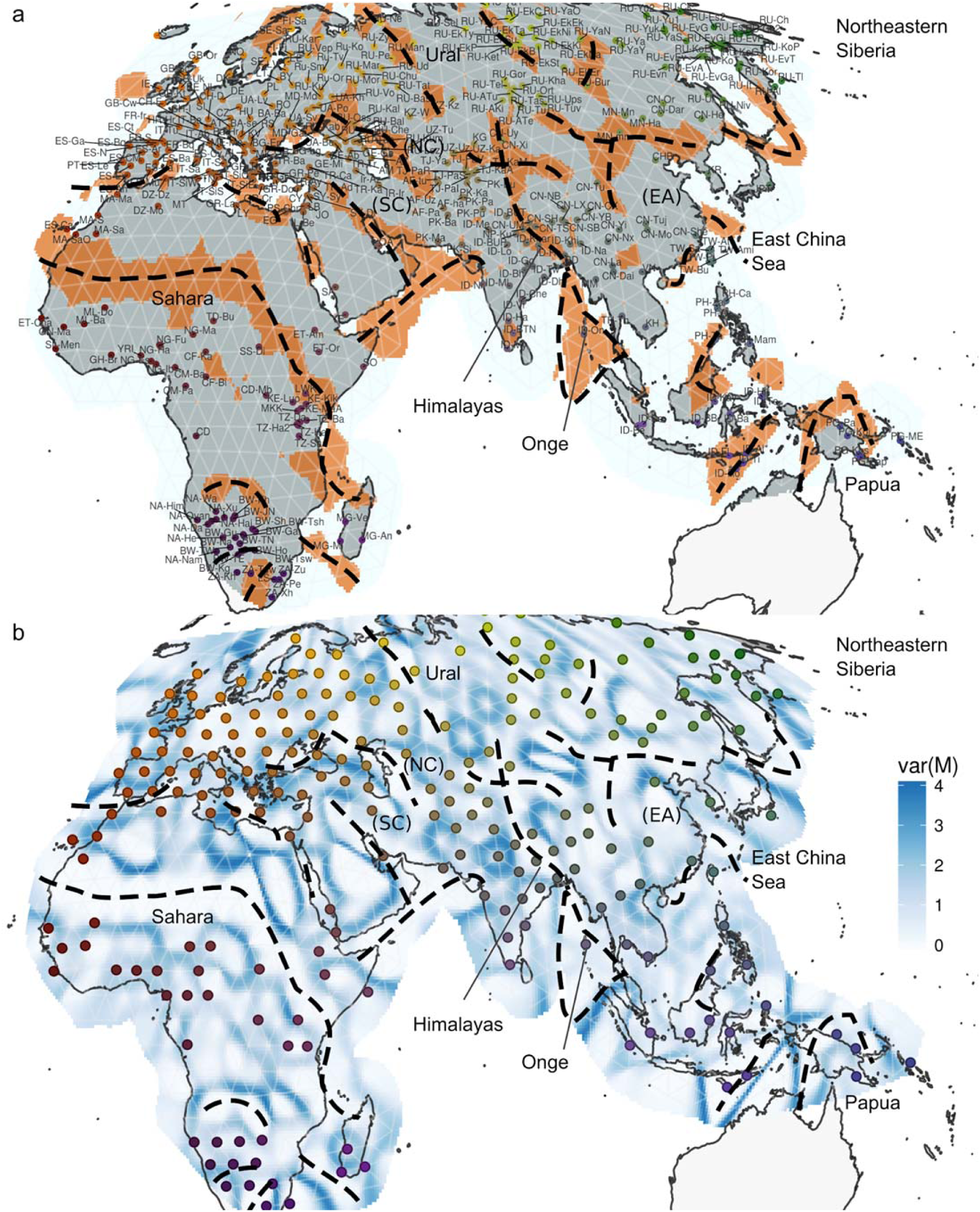
a: Location of troughs (below average migration rate in more than 95% of MCMC iterations) are given in brown. Sample locations and EEMS grid are displayed. **b:** Posterior variance on migration rate parameters. Note that most significant features are in low variance regions, but that they are often surrounded by high-variance regions, implying the exact boundary of troughs is estimated with uncertainty. Grid-fitted sample locations are displayed. Annotation in both panels is identical to Figure 1a.

**Extended Data Figure 3:**
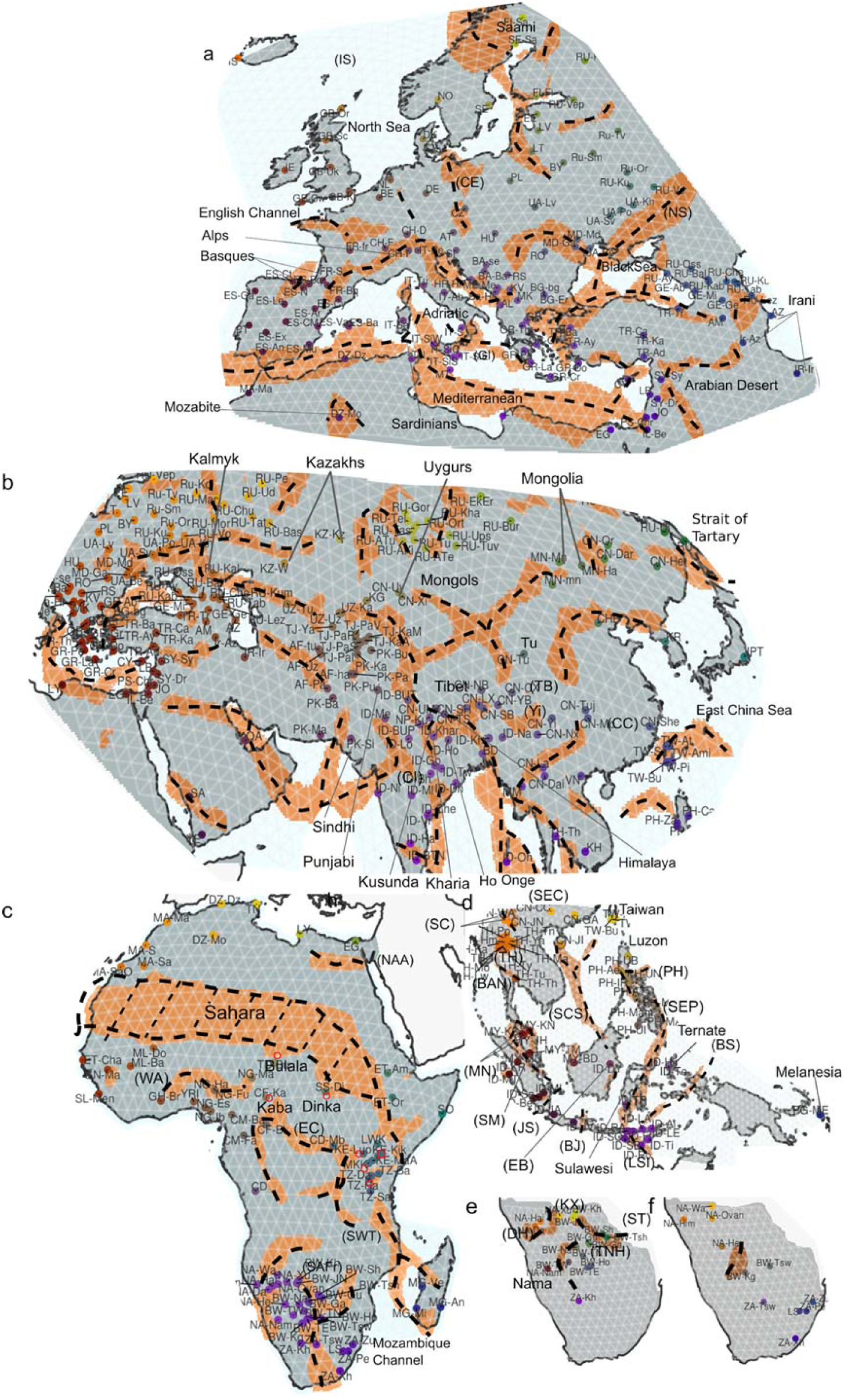
Location of troughs (below average migration rate in more than 95% of MCMC iterations) are given in brown. Sample locations and EEMS grid are displayed for **a**: WEA **b**: CEA **c**: AFR **d:** SAHG and **e**: SEA analysis panels. Annotation in all panels is identical to Figure 2.

**Extended Data Figure 4:**
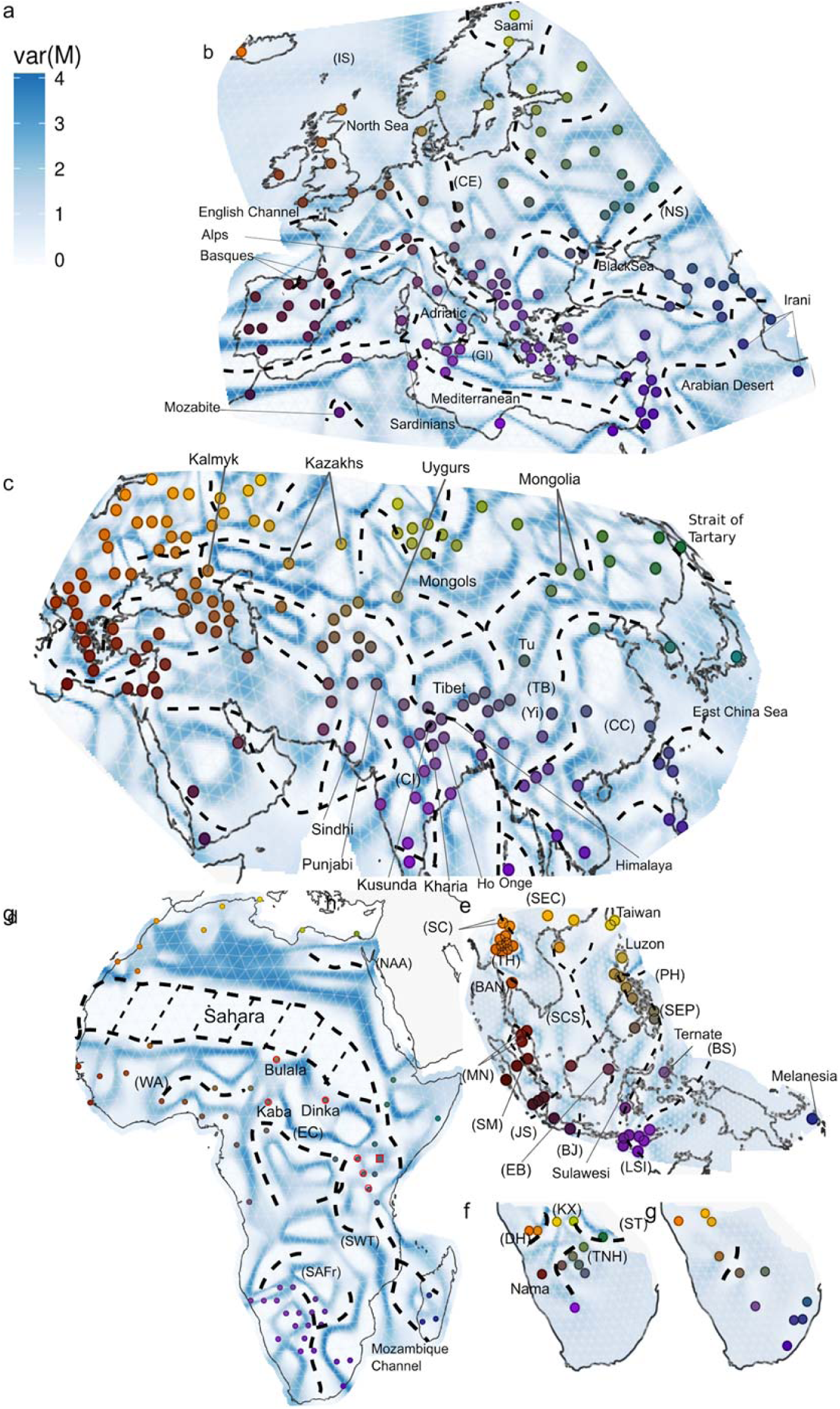
Posterior variances in migration rate parameters. Grid-fitted sample locations are displayed. **a**: scale bar **b:** WEA **c**: CEA **d**: AFR **e:** SAHG and **f**: SEA analysis panels. Note that most significant features are in low variance regions, but that they are often surrounded by high-variance regions, implying the exact boundary of troughs is estimated with uncertainty. Annotation of troughs and select features is identical to Figure 2.

**Extended Data Figure 5:**
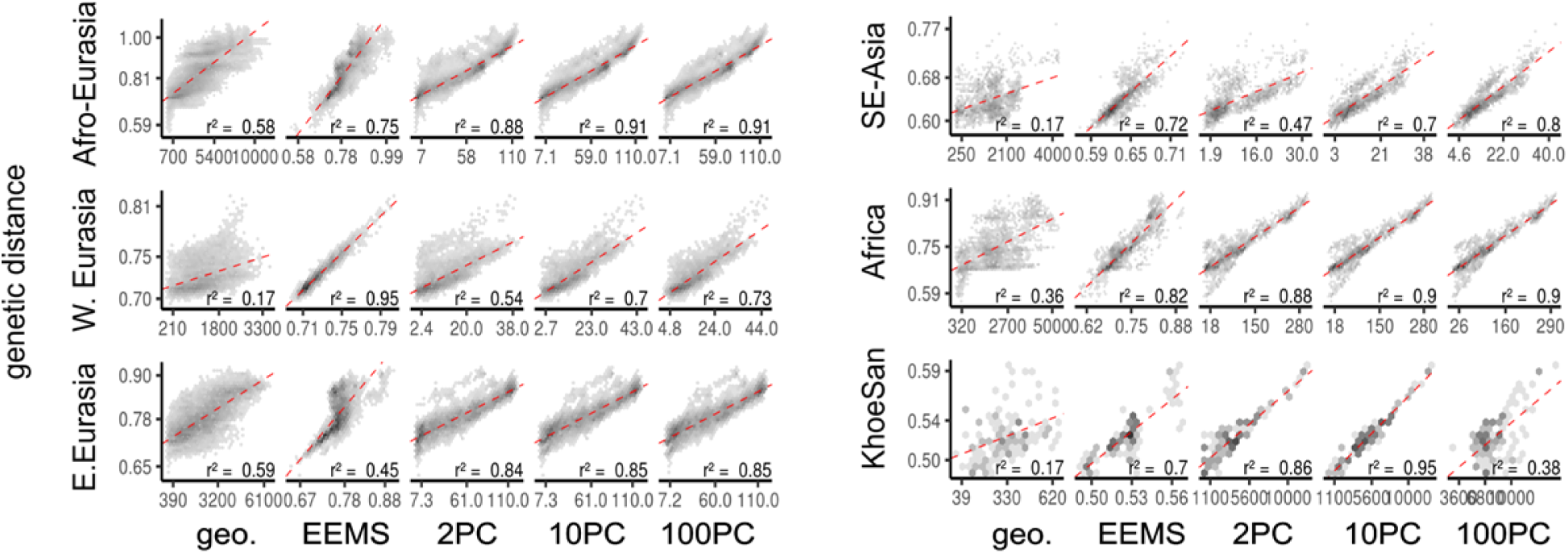
Hex-binned scatterplots of genetic distance versus geographic distance (in km), predicted distance via EEMS model fit, and predicted distance via a ten-component PCA, for all panels. Darker areas correspond to bins with more points. The fit of a simple linear regression (red dashed lines) and r^2^ are given.

**Extended Data Figure 6:**
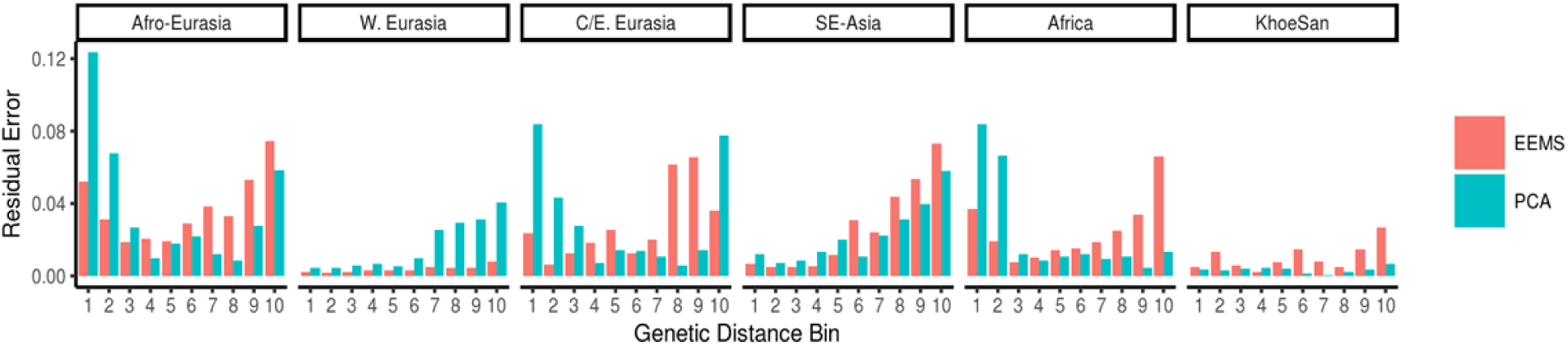
Comparing Fit of PCA and EEMS. We show the relative error of EEMS (red) and PCA(blue, first 10 PCs) for all pairs, stratified by genetic distance. For each panel, all pairwise genetic distances were distributed in ten bins of equal size, for which we then computed the median absolute error of the fitted model vs the observed distances. For W. Eurasia and SE-Asia, EEMS fits uniformly better than PCA. In the Afro-Eurasian, Central/Eastern Eurasian and African panel, EEMS fitts better for smaller distances, but the fit is worse for larger distances. For the KhoeSan, EEMS fits worse than PCA for all distance bins.

**Extended Data Figure 7:**
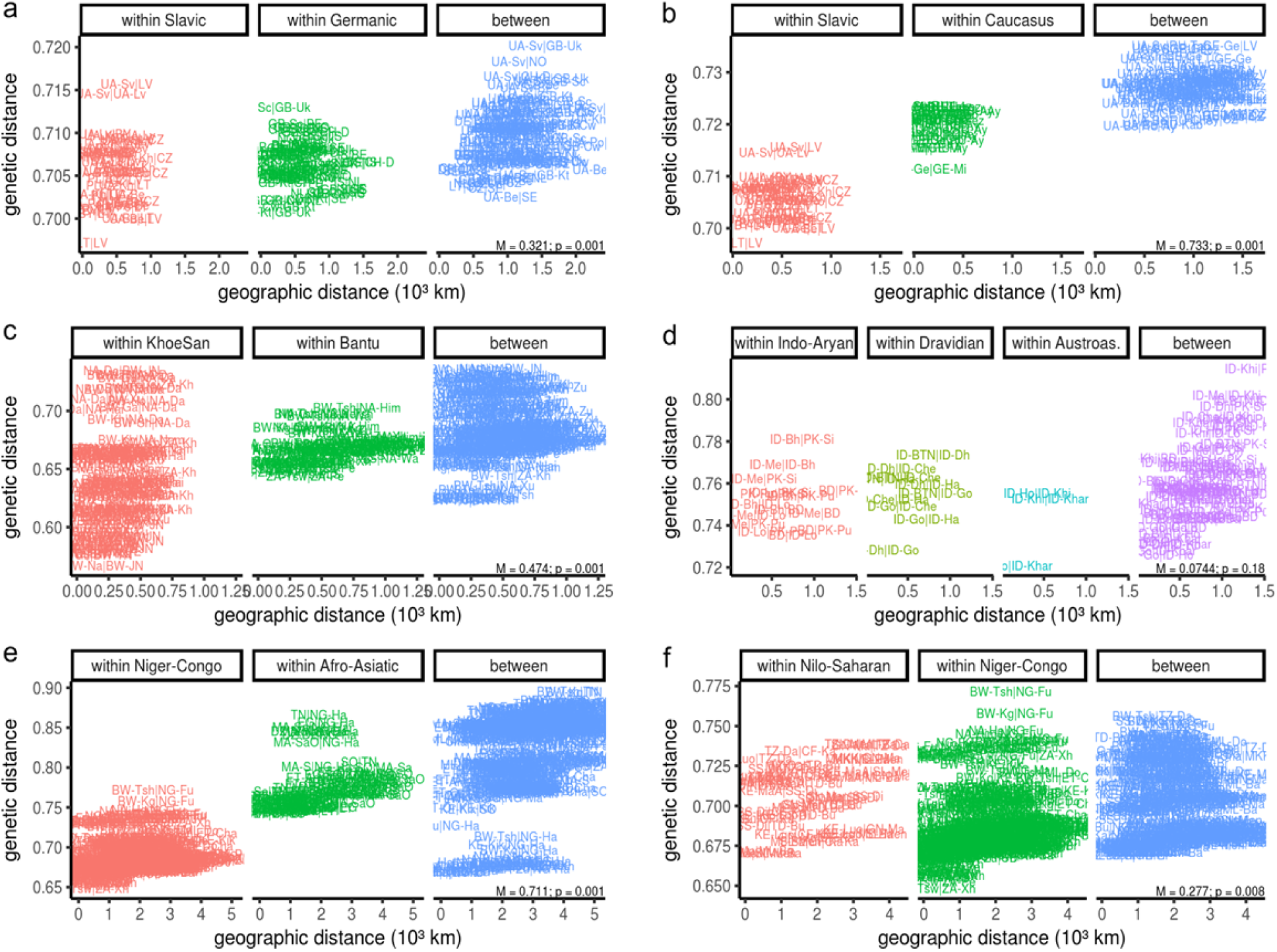
Genetic vs. geographic distance within and between language groups. The eems-plots revealed several troughs aligning with differences in linguistic groups. We show the pairwise relationship of genetic and geographic differences within- and between adjacent language groups mentioned in the main text for a. Slavic and Germanic speakers (WEA panel) b. Slavic and Caucasus languages (WEA), c. KhoeSan and Bantu languages (Southern Africa) d. Indo-Aryan, Dravidian and Austroasiatic (CEA) e. Niger-Congo and Afro Asiatic (AFR) and f. Nilo-Saharan and Niger-Congo (AFR).

**Extended Data Figure 8:**
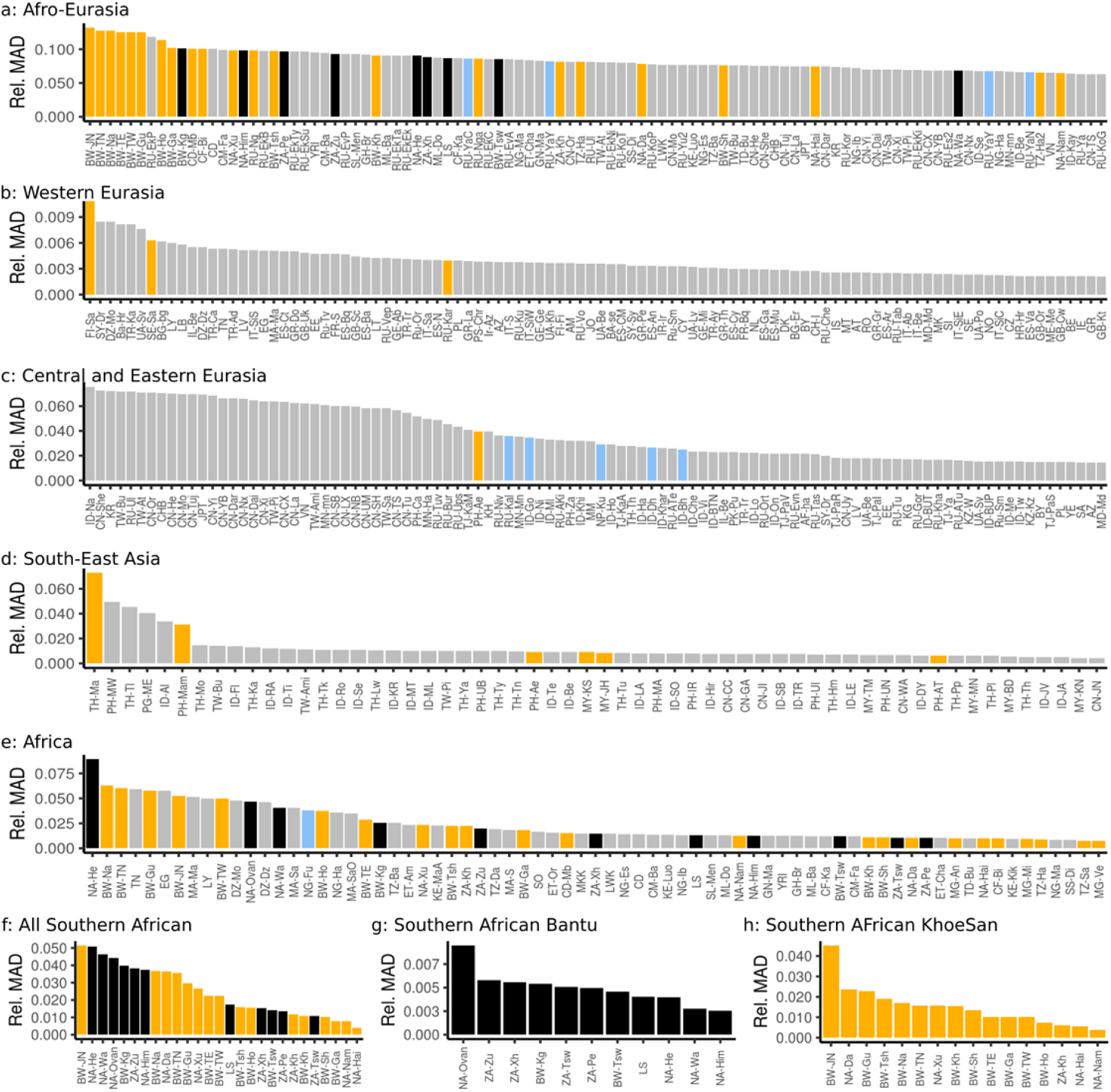
EEMS-fit residuals. For each population, we show the median absolute deviation (MAD) of the observed vs EEMS-fitted genetic distances, normalized by the median distance for this population. yellow: Hunter-Gatherers; Black: Southern African Bantu speakers; Blue: Populations with a recent admixture or displacement.

**Extended Data Figure 9:**
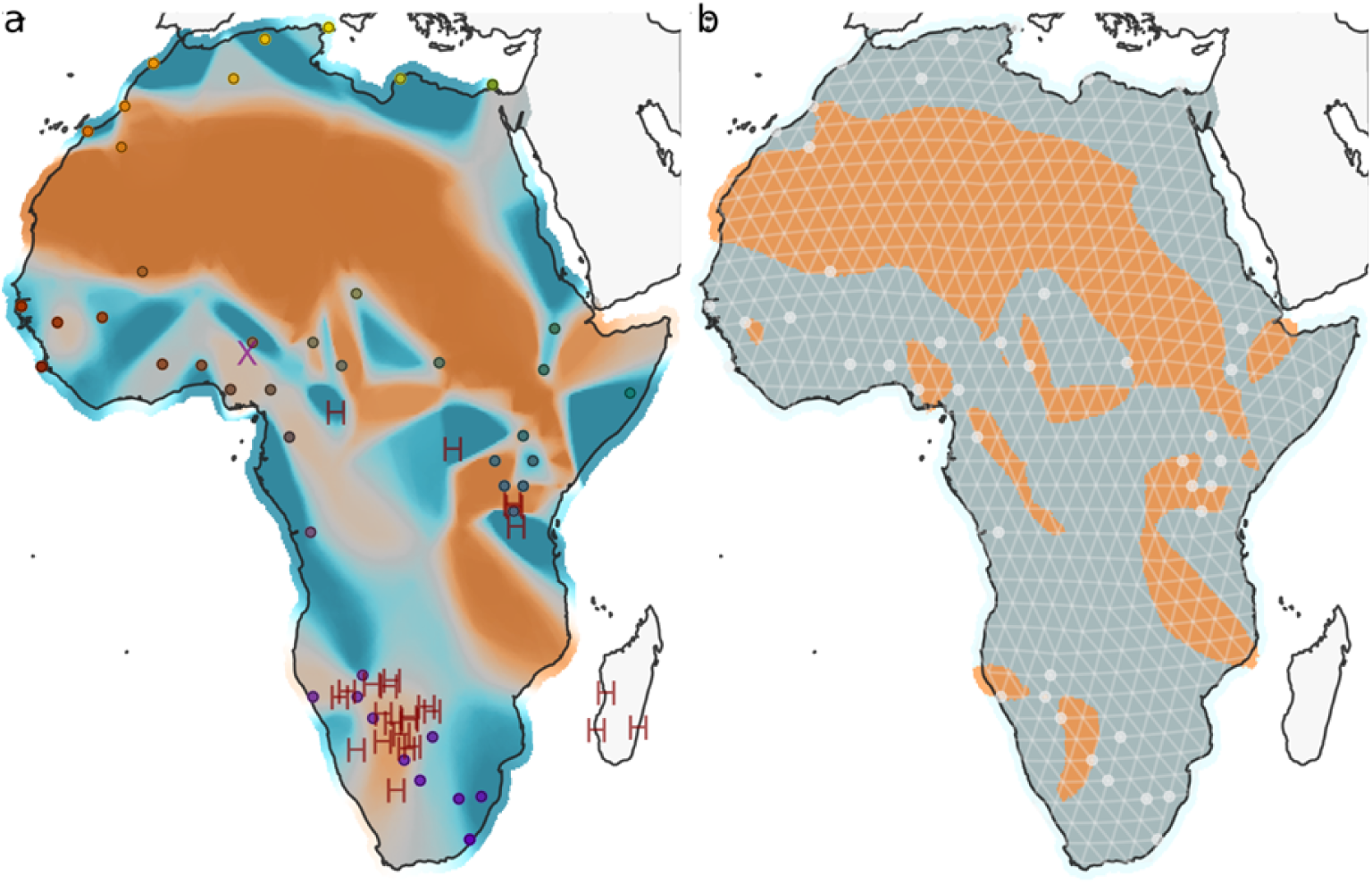
Alternative Africa analysis. To assess the effect of populations that may not be modelled well by EEMS (admixed or hunter-gatherer populations), we provide supplemental analyses of Africa with several populations excluded from the model fit. **a:** EEMS-map and **b:** location of troughs for Africa. Excluded populations are annotated with H (Hunter-gatherers) and X (admixed). With this filtering (in particular removing the Hadza and Sandawe), the Eastern African trough between Afro-Asiatic speakers and Nilo-Saharan / Niger-Congo speakers (seen in Figures 1 and 2g) vanishes.

